# Bionic 3D printed corals

**DOI:** 10.1101/834051

**Authors:** Daniel Wangpraseurt, Shangting You, Farooq Azam, Gianni Jacucci, Olga Gaidarenko, Mark Hildebrand, Michael Kühl, Alison G. Smith, Matthew P. Davey, Alyssa Smith, Dimitri D. Deheyn, Shaochen Chen, Silvia Vignolini

## Abstract

Symbiotic corals have evolved as a highly optimised photon augmentation system leading to space-efficient microalgal growth and photosynthetic quantum efficiencies that approach theoretical limits^1–3^. Corals are characterized by an elastic animal tissue hosting microalgae and a light scattering calcium carbonate skeleton that maximizes light delivery towards otherwise shaded algal-containing tissues^4,5^. Rapid light attenuation due to algal self-shading is a key limiting factor for the upscaling of microalgal cultivation^6,7^. Coral-inspired light management systems could overcome this limitation and facilitate scalable bioenergy and bioproduct generation^8,9^. Here, we developed 3D printed bionic corals capable of growing various types of microalgae with cell densities approaching 10^9^ cells mL^-1^, up to 100 times greater than in liquid culture. The hybrid photosynthetic biomaterials are produced with a new 3D bioprinting platform which mimics morphological features of living coral tissue and the underlying skeleton with micron resolution, including their optical and mechanical properties. The programmable synthetic microenvironment thus allows for replicating both structural and functional traits of the coral-algal symbiosis. Our work defines a new class of bionic materials capable of interacting with living organisms, that can be exploited for the design of next generation photobioreactors^7^ and disruptive approaches for coral reef conservation^10^.

---

Our bioprinting platform is capable of 3D printing optically-tunable photosynthetic matter that mimics coral tissue and skeleton morphology with micron-scale precision (Fig. 1a-g). In principle, our technique allows replication of any coral architecture (Extended Data Fig. 1), providing a variety of design solutions for augmenting light propagation. Fast-growing corals of the family *Pocilloporidae* are particularly relevant for studying light management. Despite high algal cell densities in their tissues (1 × 10^6^ cells per cm^2^ surface area), the internal fluence rate distribution is homogenous, avoiding self-shading of the symbiotic microalgae^11^. The photon distribution is mainly managed by the aragonite skeleton, where light leaks out of the skeleton and into coral tissue, supplying photons deep within the corallite^4,12^. Additionally, light can enter the coral tissue more easily than it can escape, as low angle upwelling light is trapped by refractive index mismatches between the coral tissue and the surrounding seawater^13^. We therefore mimicked these light management strategies and designed a bionic coral for enhanced microalgal light absorption and growth.

**Fig. 1.**
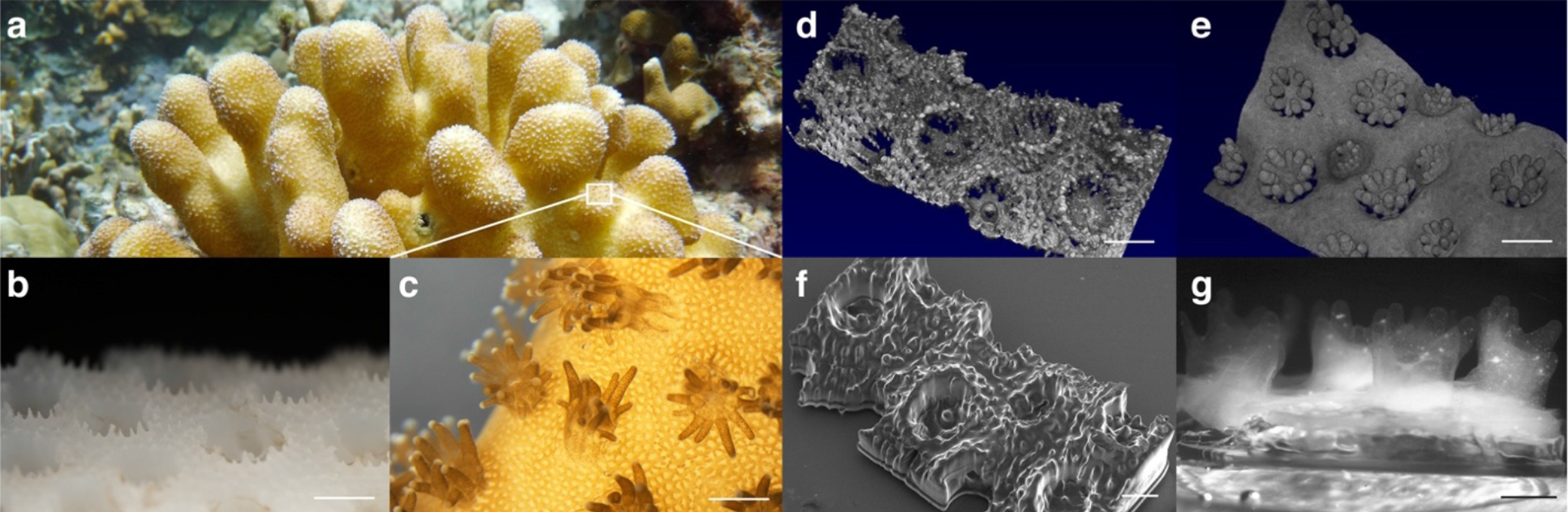
Structure of natural and 3D printed bionic corals. Colony of the coral *Stylophora pistilla* growing at about 10 m depth on Watakobi Reef, East Sulawesi, Indonesia (**a**). Close-up photograph (**b, c)** and optical coherence tomography scanning (**d, e)** of coral skeleton and coral tissue, respectively. Scanning electron microscopy image of successful 3D printed skeleton replica showing corallites in 1:1 scale relative to the original design (**f**). Photograph of living bionic coral growing *Symbiodinium sp*. microalgae (**g**). The living tissue was printed on top of the skeleton mimic and the bionic coral was cultured for 7 days. Scale bar = 1 mm (**b-f**) and 750 μm (**g**).

To precisely control the scattering properties of the bio-inspired artificial tissue and skeleton, we developed a 2-step continuous light projection-based approach for multilayer 3D bioprinting (Methods). Optimization of the printing approach required a delicate balance between several parameters including printability (resolution and mechanical support), cell survival, and optical performance (Methods). The artificial coral tissue constructs were fabricated with a novel bio-ink solution, in which the symbiotic microalgae (*Symbiodinium sp*.) were mixed with a photopolymerizable gelatin-methacrylate (GelMA) hydrogel and cellulose-derived nanocrystals (CNC), the latter providing mechanical stability and allowed tuning of the tissue scattering properties^14,15^. Similarly, the artificial skeleton was 3D printed with a polyethylene glycol diacrylate-based polymer (PEGDA)^16^ doped with CNC.

Based on optimization via experiments and optical simulations (Fig. 2), the functional unit of the artificial skeleton was an abiotic cup structure, shaped like the inorganic calcium carbonate corallite (1 mm in diameter and depth) and tuned to redistribute photons via broadband diffuse light scattering (scattering mean free path = 3 mm between 450-650 nm) and a near isotropic angular distribution of scattered light (Fig. 2h, Extended data Fig. 2), similar to the optical properties of the skeleton of fast growing intact corals^5,12^. The coral-inspired tissue had cylinder-like constructs (200 μm wide and 1mm long) radially arranged along the periphery of the corallites mimicking coral tentacles, which serve to enhance surface area exposed to light^17^ (Fig. 2a). We designed the bionic coral tissue to have a forward scattering cone (Fig. 2h), which enabled light to reach the diffusely backscattering skeleton (Fig. 2a).

**Fig. 2.**
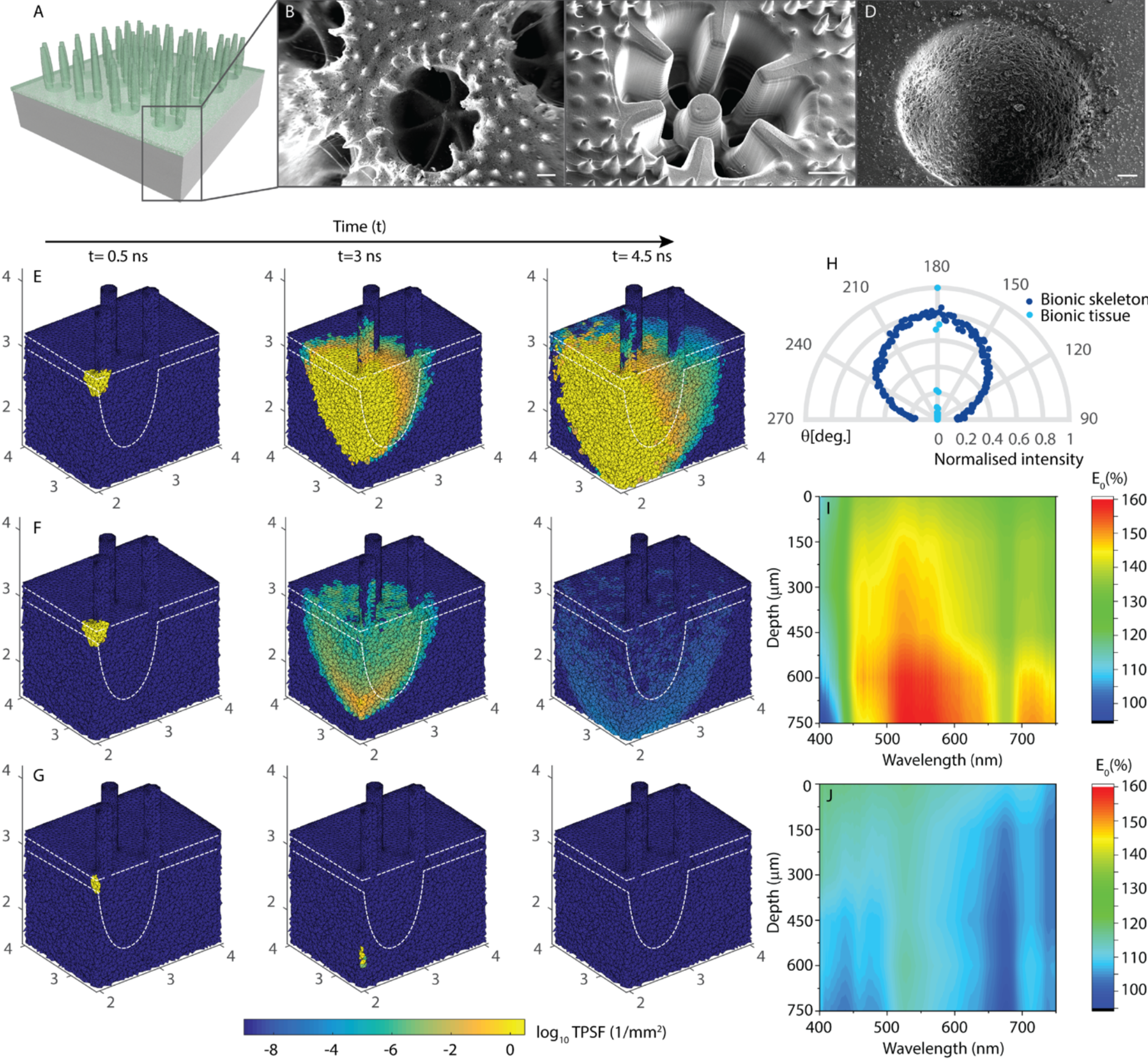
Optical properties of 3D printed bionic coral tissue and skeleton. 3D rendering of final bionic coral design (**a**). Bionic skeletal design optimization (**b-d**) showing SEM images of the original *Stylophora pistillata* corallite architecture (scale bar = 200μm) (**b**), a 3D printed intermediate skeleton design (scale bar = 300 μm) (**c**) and the final bionic skeleton doped with CNC aggregates (scale bar = 100 μm) (**d**). 3D tetrahedral mesh-based Monte Carlo simulation (**e-g**). Light (675 nm) is irradiated over the connecting tissue (red arrow) as a collimated pencil beam. The time-resolved solution of photon migration (temporal point spread function, TPSF [1/mm^2^]) is shown after 0.5 ns (left column), 3 ns (center column), and 4.5 ns (right column) in a cross-cut view of the 2-layer bionic coral (**e**), a 1-layer bionic tissue (**f**) and non-scattering GelMA (**g**). The microalgal density in the tissue component is identical for all simulations (μ_a_=15 mm^-1^). The angular distribution of forward scattered light (q=270°–90°) at 550 nm is shown as normalized transmittance (**h**). Microprobe-based fluence rate measurements (E0 in % of incident irradiance) for the bionic coral (**i**) and a flat slab of GelMA (**j**) both with a microalgal density of 5.0 × 10^6^ cells mL^-1^.

Our bionic coral increased the photon residence time as light travelled through the algal culture (Fig. 2e), consequently enhancing the chance of light absorption for photosynthesis by algae deeper within it^13^. As photons travelled through the bionic skeletal cup, they were redirected into the photosynthetic tissue and the contribution of such scattered light increased with depth, effectively delivering light to the deepest part of the bionic coral (Fig. 2i). This photon augmentation strategy led to a steady increase of irradiance with tissue depth, which counter balanced the exponential light attenuation by photopigment absorption (Fig. 2g)^18^. Compared to a flat slab of biopolymer (GelMA) with the same microalgal density (5.0 × 10^6^ cells mL^-1^), the fluence rate (for 600 nm light) measured in the photosynthetic layer of the bionic coral was more than 1.5-fold enhanced at 750 μm depth (Fig. 2i,j).

In order to evaluate the growth of a commercially-relevant microalgal species in our bionic coral, we cultured the green alga *Marinichlorella kaistiae* KAS603^19^ (Fig. 3a-d, Methods). Although the main focus of our bionic coral design was to improve light management, it also allows for growing microalgae without the need for energy-intensive turbulent flow and mixing, which is otherwise required for optimal nutrient and light delivery in photobioreactors^6^. This is accomplished by the combination of the bionic tissue and skeleton replica morphology, the tissue mechanical properties (average Young’s modulus, *E* = 4.3 kPa, Extended Data Fig. 4) and its porosity (pore size diameter = 5-40 μm) (Methods). We grew *M. kaistiae* KAS603 under no-flow conditions and low incident irradiance (Ed = 80 μmol photons m^-2^ s^-1^) in our bionic coral, where it reached algal cell densities of >8 × 10^8^ cells mL^-1^ by day 12 (Fig. 3a). This is about one order of magnitude higher than the maximal cell densities reported for this algal species when grown in flasks under continuous stirring^19^. Despite such high algal cell densities, irradiance did not limit growth at depth, and about 80% of the incident irradiance remained at 1 mm depth within the bionic coral tissue construct (Fig. 3b). In comparison, biofilm-based photobioreactors show exponential light attenuation characterized by a virtual depletion of irradiance within 200-300 μm of the biofilm thickness^20^. We observed that *M. kaistiae* KAS603 grew in the bionic tissue as dense aggregates (sphericity 0.75 ± 0.09 SD, diameter = 30-50 μm; Fig. 4a-d). Algal photosynthesis within the tissue construct yielded a net photosynthetic O_2_ production of 0.25 nmol O_2_ cm^-2^ s^-1^ at the polyp tissue surface (Fig. 3c). Gross photosynthesis within 8-day old bionic coral polyps was enhanced at a depth of 300 μm compared to gross photosynthesis rates measured at the surface of the bionic coral tissue (Fig. 3d).

**Fig. 3.**
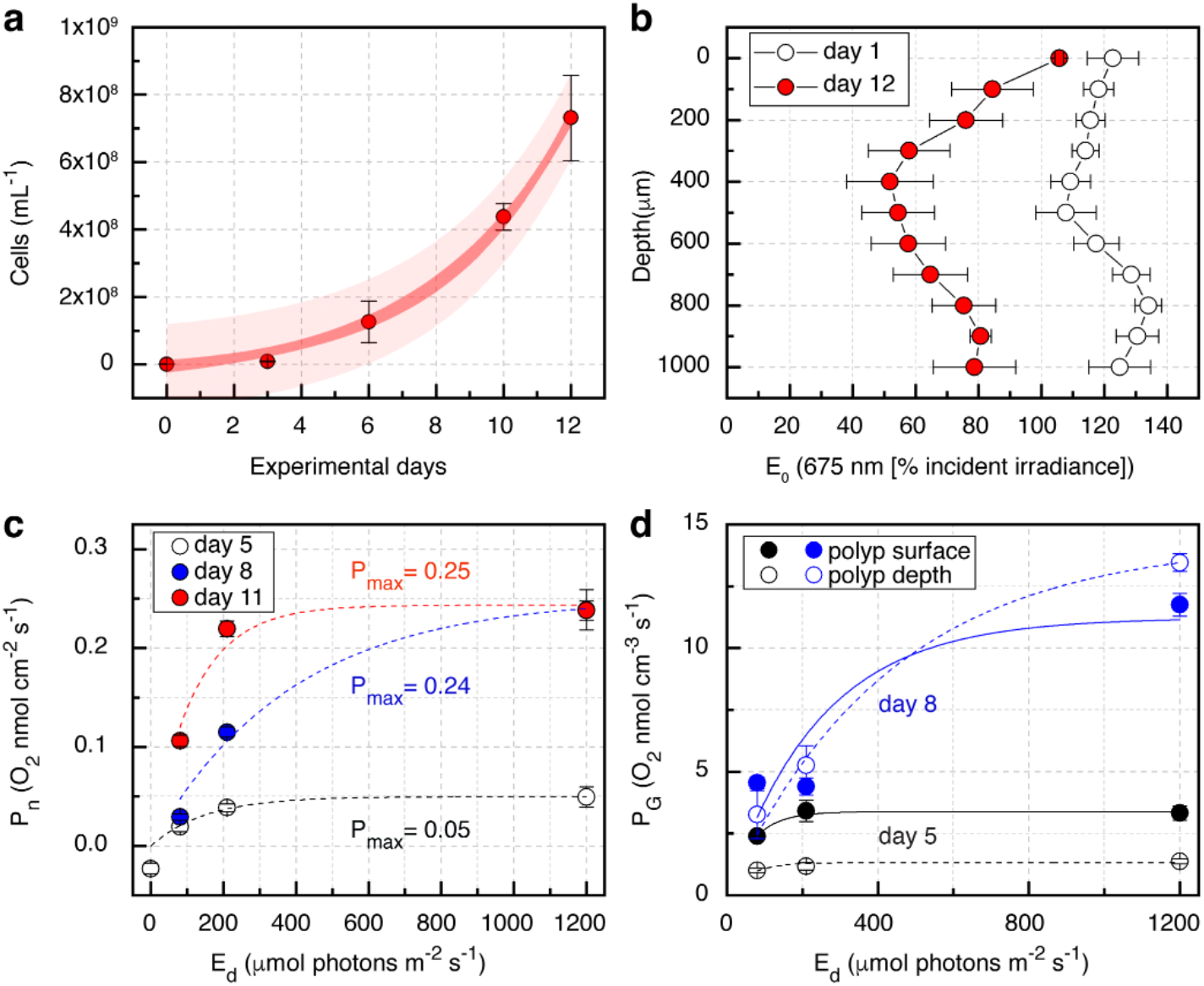
Performance testing of 3D printed bionic coral. Growth of *Marinichlorella kaistiae* KAS603 in bionic coral (**a**). Data are means (± SEM, *n* = 3-6 bionic coral prints). Dark and bright red areas show 95% confidence and prediction intervals, respectively. Vertical attenuation of fluence rate (E0 at 675 nm) at the beginning (day 1) and end of the performance test (day 12) (**b**). Net photosynthetic rates at day 5, 8 and 11 (**c**). Lines represent curve fits (see Methods). Gross photosynthetic rates at day 5 (black) and day 8 (blue) (**d**). Measurements were performed with O_2_ microsensors at the center of the corallite cup surface (closed symbols/solid lines) and at a vertical depth of 300 μm (open symbols/dashed lines). Symbols are means (± SEM, *n* = 3-6 bionic coral prints), lines are curve fits.

**Fig. 4.**
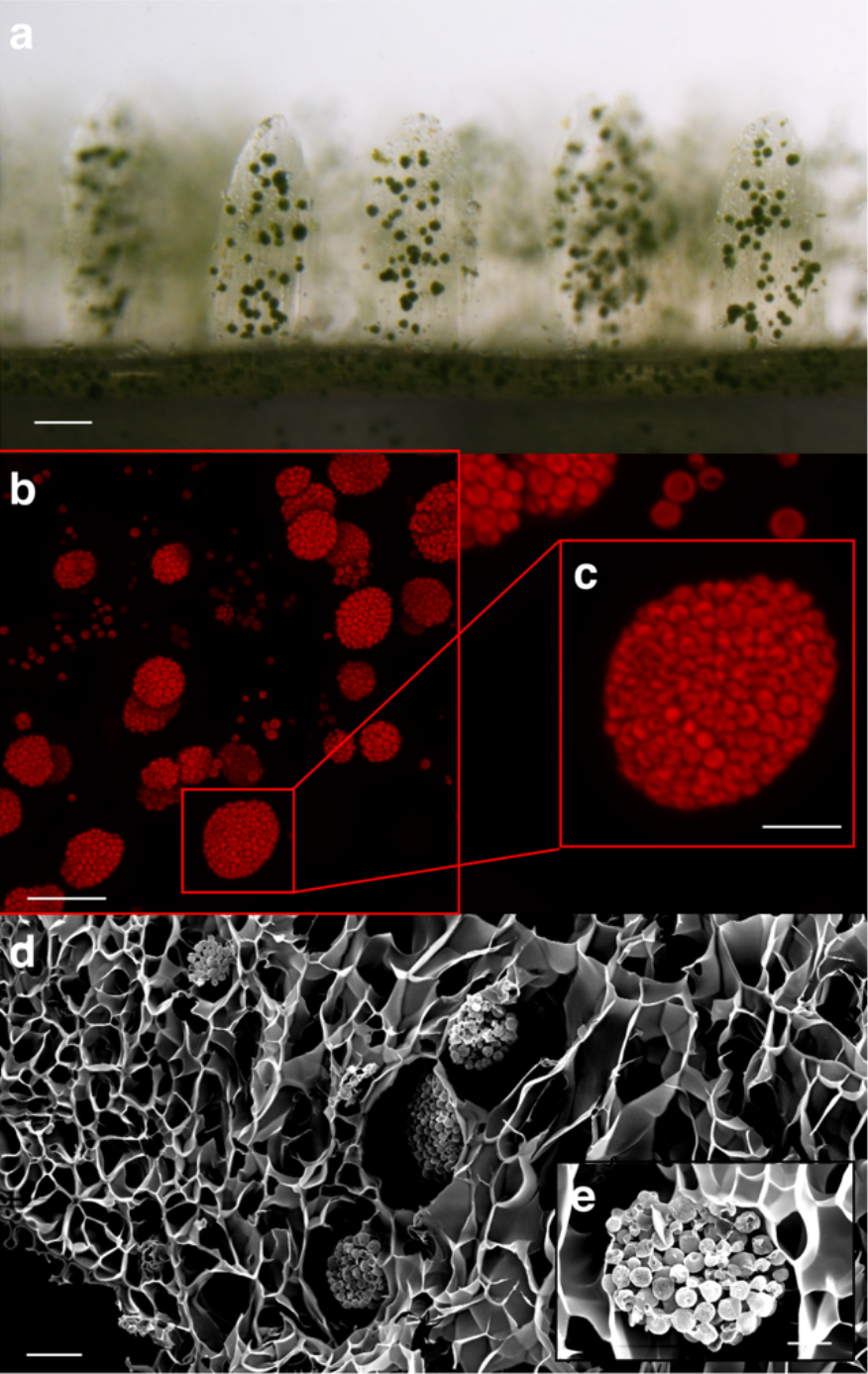
Living 3D printed bionic coral. Horizontal view of 7-day old bioprinted construct, showing aggregates of the green microalga *Marinichlorella kaistiae* KAS603 (scale bar = 100 μm) (**a).** Maximum *z* projection of confocal images showing chlorophyll *a* fluorescence of bionic tentacles (scale bar = 50 μm) (**b**) and a *M. kaistiae KAS603* aggregate (scale bar = 20 μm) (**c**). SEM image of bionic tissue showing porous tissue scaffolds (scale bar = 20 μm) (**d**) and a close-up of a microalgal aggregate (scale bar = 10 μm) (**e**).

Bionic corals enable microalgal cultivation with strongly reduced energy maintenance requirements, as they do not require water mixing and the fluence rate within the biomaterial is up to two-fold enhanced relative to the incident light source (Methods, Fig. 2i,j). The light redistribution in the bionic corals can be controlled by adjusting the concentration and dimensions of the CNCs in the biopolymer (Extended Data Fig. 2). This approach allows the design of various coral host mimics (Extended Data Fig. 1) accommodating algal strains with different photon requirements for optimal productivity^21^. Compared to commercial outdoor raceway ponds that achieve an aerial productivity of ~4-20 g m^-2^ day^-1(*21,22*)^, our bionic corals, if scaled appropriately, would use up to 150 times lower volume and achieve approximately 2.55 g m^-2^ day^-1^ (Methods). The high spatial efficiency of the bionic coral system is therefore particularly suitable for the design of compact photobioreactors for algal growth in dense urban areas or as life support systems for space travel^18,23^. Moreover, bionic corals allow investigation of the cellular activity of specific *Symbiodinium* strains, while mimicking the natural optical and mechanical microenvironment of different corals, providing an invaluable tool for advancing the frontiers of animal-algal symbiosis and coral bleaching research. We therefore anticipate that bionic corals will find multidisciplinary applications from biological studies to commercial technologies for efficient photon augmentation for bioenergy and bioproducts.

## METHODS

### Optical coherence tomography imaging

To create a digital mask of natural coral surfaces, a spectral-domain (SD) optical coherence tomography (OCT) system (Ganymede II, Thorlabs GmbH, Dachau, Germany) was used to image living corals (Extended Data Figure 1). The OCT system was equipped with a superluminescent diode (centered at 930 nm) and an objective lens (effective focal length = 36 mm) (LSM03; Thorlabs GmbH, Dachau, Germany) yielding a z-resolution of 4.5 μm and a x-y resolution of 8 μm in water. The imaged coral species (*Pavona cactus, Stylophora pistillata, Pocillopora damicornis, Favites flexuosa*) were maintained at the Centre Scientifique de Monaco, and corals were imaged under controlled flow and irradiance conditions. For OCT imaging of bare coral skeletons, the living tissue was air brushed off the skeleton. The skeleton was carefully cleaned before imaging the bare skeleton in water. OCT scanning was performed as described previously^17^.

### Surface rendering of OCT data

OCT data was extracted as multiple 16-bit TIFF image stacks and imported into MATLAB (Matlab 2018a). Image acquisition noise was removed via 3D median filtering. Segmentation of the outer tissue or skeletal surface was done via multilevel image thresholding using Otsu’s method on each image of every TIFF stack. The binary files were exported as *x,y,z* point clouds and converted to a *stl* file format, which could be sliced into 2D image sequences for bioprinting^24^. If the generated *stl* files showed holes in the surface mesh, these holes were manually filled using *Meshlab* (Meshlab 2016).

### Algal-biopolymer design optimization

Key characteristics to achieve in material design were 1) high microalgal cell viability and growth, 2) microscale printing resolution, and 3) optimization of light scattering and biomechanical parameters including material stiffness, porosity and molecular diffusion. The photo-induced, free radical polymerization mechanism underlying our 3D printing technique allowed us to precisely control the mechanical properties via modulating the crosslinking density of the polymerized parts^*27*^. Any material and fabrication parameters (e.g., light intensity, exposure time, photoinitiator concentration, material composition) that affect the crosslinking density can be employed to tune the mechanical properties of the printed parts. Initially, different concentrations of prepolymer and photoinitiator combinations were evaluated, including glycidal methacrylate-hyaluronic acid (GM-HA), gelatin methacrylate (GelMA), polyethylene glycol diacrylate (PEGDA), and poly(lactic acid), together with the photoinitiators Irgacure 651 and lithium phenyl-2,4,6 trimethylbenzoylphosphinate (LAP). Cell viability and growth were higher in GelMA compared to PEGDA (data not shown), probably due to favorable diffusion characteristics of GelMA because of its highly porous microstructure^25^, while PEGDA has stronger mechanical stiffness. Therefore, we combined PEGDA with GelMA to make a mechanically robust and tunable hydrogel. GelMA was initially doped with graphene oxide, which enhanced mechanical stability but limited light penetration and cell growth. We developed a photopolymerization system using 405 nm light to avoid UV damage to the algae.

To optimize light scattering, we first mixed the hydrogel with different concentrations of SiO2 particles (Sigma-Aldrich, USA) that were in a size range (about 10 μm) to induce broadband white light scattering with high scattering efficiency. However, when mixed into the hydrogels, the SiO2 particle showed a vertical concentration gradient related to the particle sinking speed in the gel. Instead, we used cellulose nanocrystals (CNCs), which exhibit suitable light scattering, mechanical properties and low mass density^26^. CNCs can be considered as rod-shaped colloidal particles (typical length of 150 nm and a width of a few nm in diameter), which have high refractive index (about 1.55 in the visible range). CNCs have received an increasing interest in photonics, due to their colloidal behaviour and their ability to self-assemble into cholesteric optical films^26^. In the 3D bioprinted coral skeleton samples that contain 7% CNCs (w/v), we found that CNCs aggregated to form microparticles with a size range of 1-10 μm. These aggregated microparticles are highly efficient white light scatterers (Extended Data Fig. 2a). In contrast, the 3D bioprinted bionic coral tissue constructs contained only 0.1% CNCs (w/v), and we did not observe any aggregated CNC microparticles.

The printing polymer solution (bio-ink) for the bionic coral tissue constructs was made up of final concentrations of: *Marinichlorella kaistiae* KAS603 (1×10^6^ cells mL^-1^), GelMA (5% w/v), LAP (0.2 % w/v), food dye (1% v/v), PEGDA (6000 Da; 0.5% w/v), CNC (0.1 % w/v), and artificial seawater (ASW; 93.2 %). The yellow food dye (Wilton^®^ Candy Colors) was added to limit the penetration of polymerization-inducing light into the bio-ink. This leads to higher light absorption relative to scattering and enhanced the spatial resolution of the printing^27^. The food dye is non-toxic and diffuses out after 24 hr (data not shown).

### Polymer synthesis

PEGDA (mol wt, Mn = 6000) was purchased from Sigma–Aldrich (USA). GelMA was synthesized as described previously^28^. Briefly, porcine gelatin (Sigma Aldrich, St. Louis, MO, USA) was mixed at 10% (w/v) into ASW medium (see above) and stirred at 60°C until fully dissolved. Methacrylic anhydride (MA; Sigma) was added until a concentration of 8% (v/v) of MA was achieved. The reaction continued for 3 hr at 60°C under constant stirring. The solution was then dialyzed against distilled water using 12–14 kDa cutoff dialysis tubing (Spectrum Laboratories, Rancho Dominguez, CA, USA) for 7 days at 40°C to remove any unreacted methacrylic groups from the solution. The GelMA was lyophilized at −80°C in a freeze dryer (Freezone, Labonco) for one week to remove the solvent.

CNC suspensions were prepared as described previously^26^. The photoinitiator lithium phenyl-2,4,6 trimethylbenzoylphosphinate (LAP) was synthesized as described previously^29^. Freeze dried LAP was dissolved with ASW, and CNC was dispersed in the LAP solution via vortexing for about 5 min.

### Continuous multilayer 3D bioprinting of bionic coral

The bionic coral design was developed as an optimization between algal growth rates, optical performance and the outcome of optical models (Fig. 2, Extended Data Figure 2, 3). The final bionic coral was designed in CAD software (Autodesk 3ds Max, Autodesk, Inc, USA) and was then sliced into hundreds of digital patterns with a custom-written MATLAB program. The digital patterns were uploaded to the a digital micromirror device (DMD) in sequential order and used to selectively expose the prepolymer solution for continuous printing. A 405-nm visible LED light panel was used for photopolymerization. A digital micromirror device (DMD) consisting of 4 million micromirrors modulated the digital mask projected onto the prepolymer solution for microscale photopolymerization^27^. The continuous movement of the DMD was synchronized with the projected digital mask to create smooth 3D constructs that are rapidly fabricated without interfacial artifacts. To print the bionic coral, a 2-step printing approach was developed. In the first step, the PEGDA bio-ink was used to print the coral inspired skeleton. The resulting hydrogel was attached to a glass slide surface, washed with DI water and then dried with an air gun. In the second step, the algal cell-containing bio-ink for tissue printing was then injected with a pipette into the skeletal cavities in order to fill the air gaps. The gap-filled skeletal print was repositioned at the identical spot on the bioprinter, and the bionic coral tissue mask was loaded. The z-stage was moved such that the surface of the skeletal print touched the glass surface of the bioprinter.

### Algal stock culture maintenance

Two microalgal species were chosen for inclusion in 3D bioprinted polymers: dinoflagellates belonging to the genus *Symbiodinium* and the green alga *Marinichlorella kaistiae*. Stock cultures of *Symbiodinium* strains A08 and A01 (obtained from Mary Coffroth, University of Buffalo) were cultured in F/2 medium in a 12h/12h light:dark cycle under an irradiance (400-700 nm) of 200 μmol photons m^-2^ s^-1^. Wild type *M. kaistiae* strain KAS603^19^ were obtained from Kuehnle AgroSystems, Inc. (Hawaii) and were cultivated at 25°C in artificial seawater (ASW) medium^30^ under continuous light from cool white fluorescent lamps (80 μmol photons m^-2^ s^-1^). Stock cultures were harvested during exponential growth phase for use in bioprinting.

### Culturing of bionic coral

Bionic corals harboring *Symbiodinium sp*. or *M. kaistiae* KAS603 were cultured under similar conditions as the respective algal stock cultures (see above). Prior to bioprinting, the bioink for printing bionic coral tissue constructs was inoculated with cell densities of 1 × 10^6^ cells mL^-1^ from exponentially growing cultures. We performed growth experiments with 35 bionic corals harbouring *M. kaistiae* KAS603. The bionic corals were transferred to 6-well plates filled with 3 mL of ASW medium^30^ containing broadband antibiotics (penicillin/streptomycin, Gibco) at a concentration of 1:1000. All prints were illuminated with an incident downwelling irradiance (400-700 nm) of 80 μmol photons m^-2^ s^-1^ provided by LED light panels (AL-H36DS, Ray2, Finnex) emitting white light. The prints were grown without mixing at 25°C. The ambient growth medium was replenished at day 5 and day 10. Degradation of GelMA-based tissue occurred after about 10-14 days when bacterial abundance was kept low via antibiotic treatment. Such degradation kinetics can be advantageous for more easy harvesting of the highly concentrated microalgae that are contained within the hard PEGDA-based skeleton.

### Optical characterization of bionic coral

The angular distribution of transmitted light was measured using an optical goniometer^31^. The samples were illuminated using a Xenon lamp (Ocean Optics, HPX-2000) coupled into an optical fiber (Thorlabs FC-UV100-2-SR). The illumination angle was fixed at normal incidence and the angular distribution of intensity was acquired by rotating the detector arm with an angular resolution of 1°. To detect the signal, a 600 μm core optical glass fiber (Thorlabs FC-UV600-2-SR) connected to a spectrometer (Avantes HS2048) was used. To characterize the optical properties, the total transmitted light was measured for different sample thicknesses using an integrating sphere^31^. The samples were illuminated by a Xenon lamp (Ocean Optics, HPX-2000) coupled into an optical fiber (Thorlabs FC-UV100-2-SR), and the transmitted light was collected with an integrating sphere (Labsphere Inc.) connected to a spectrometer (Avantes HS2048). In the case of the skeleton-inspired samples, where the light is scattered multiple times before being transmitted, the light transport can be described by the so-called diffusion approximation^32^. In this regime, the analytical expression, which describes how the total transmission (*T*) scales with the thickness (*L*) for a slab geometry, is given as^32^:

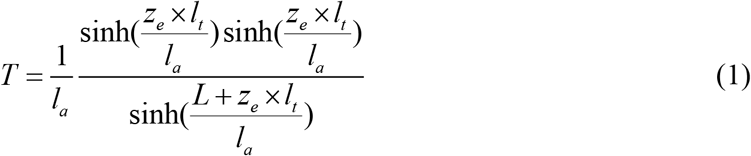

where *l_a_, l_t_* and *z_e_* are the absorption length, the transport mean free path and the extrapolation length, respectively ^33^. Here, *z_e_* quantifies the effect of internal reflections at the interfaces of the sample in the estimation of *l_a_* and *l_t_*^33^. We quantified *z_e_* by measuring the angular distribution of transmitted light ^31^, *P*(*μ*), which is related to *z_e_* by the following equation^34^:

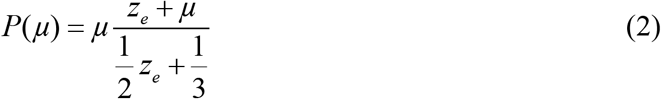

where *μ* is the cosine of the transmission angle with respect to the incident ballistic beam. The theoretical fit is shown in Figure S2C and led to a value of *z_e_* = (1.32±0.12). Once the extrapolation length was estimated, the values of *l_a_* and *l_t_* could be calculated with Eq. (1) (Extended Data Figure 2d,e). This was done with an iteration procedure to check the stability of the fit, as described previously^35^. In the bionic coral tissue, the scattering strength of the material is too low and the diffusion approximation cannot be applied. In this regime, the extinction coefficient can be estimated using the Beer-Lambert law (Extended Data Figure 2f).

The refractive index (*n*) of the bioprinted bionic coral tissue was determined with the optical goniometer to characterize the Brewster angle (*θ_B_*). A half circle of the material was printed with a diameter of 2 cm and a thickness of z = 5 mm. The Brewster angle was calculated according to Snell’s law:

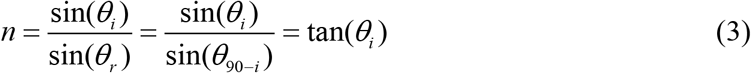

and Brewster’s law:

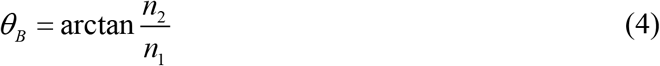

where *θ_i_* is the angle of incidence, and *θ_r_* is the angle of refraction. *n_1_* and *n_2_* are the refractive indices of the medium and the surrounding medium, respectively. For the coral-inspired tissue *q_B_* ranged between 54.0° and 55.0° yielding a refractive index of *n* = 1.37-1.40.

### 3D Monte Carlo time-of-flight photon propagation modeling

Tetrahedral meshes were generated via Delaunay triangulation using the MATLAB based program *Iso2mesh* that calls *cgalmesh*^36^. Meshing was performed with different mesh properties varying maximal tetrahedral element volume and Delaunay sphere size in order to optimize simulation efficiency. Settings were optimized for a Delaunay sphere of 1 (10 μm) and a tetrahedral element volume of 5 (50 μm). Generated tetrahedral meshes were used as source architecture for a mesh-based 3D Monte-Carlo light transport simulation (*mmclab*)^37^. The model uses the generated tetrahedral mesh and calculates photon propagation based on the inherent optical parameters, the absorption coefficient *μa* [mm^-1^], the scattering coefficient *μ_s_* [mm^-1^], the anisotropy of scattering *g* [dimensionless] and the refractive index *n* [dimensionless]^38^. The optical parameters were extracted via integrating sphere measurements (see above) and were used to calculate time-of-flight photon propagation in the bionic coral. The probe illumination was a collimated point source with varying source positions.

### Mechanical properties of bionic tissue

The Young’s modulus of the bionic coral tissue was evaluated with a microscale mechanical strength tester (Microsquisher, CellScale). Each sample was preconditioned by compressing at 4 μm s^-1^ to remove hysteresis caused by internal friction. The compression test was conducted at 10% strain with a 2 μm s^-1^ strain rate. Cylindrical constructs were 3D printed using the same bio-ink as used to print bionic coral tissue. The Young’s modulus was calculated from the linear region of the stress–strain curve^27^. Three samples were tested, and each sample was compressed three times.

### Cell counts and productivity estimates

Cell density was determined at the beginning of the experiment (day 0) and then at day 3, day 6, day 10 and day 12 of the growth experiments. To determine cell density, the construct was removed from the growth medium, and any remaining solution attached to the construct was removed with a Kimwipe. Each construct was transferred to a 1.5 mL microfuge tube and the hydrogel was dissolved via adding 600 μL trypsin solution (0.25% Trypsin/EDTA) under incubation at 37 °C for 40 min. This procedure removed the microalgal cells from the matrix allowing for cell counting via a haemocytometer^24^. The accuracy of this approach was verified by printing known cell densities (from liquid culture) and comparing it to the trypsin-based estimates yielding a deviation of < 3%. To additionally compare our cell density estimates with ash free dry weight (AFDW) of algal cell biomass [g], which is a commonly used metric in biofuels research, we determined AFDW using methods described previously^39^. AFDW was on average 3.47 × 10^-11^ g cell^-1^ (± 4.6 × 10^-13^ SE). The maximal growth rate was obtained from readings of Day10 and Day12, yielding 1.47 × 10^11^ cells L^-1^ day^-1^ or 5.1 g L^-1^ day^-1^. The aerial productivity was extrapolated to g m^-2^ day^-1^ by accounting for the area occupied by one bionic coral (6mm in length and width) and the measured productivity per bionic coral. Raceway ponds typically have a depth of 20-30 cm compared to 2 mm thickness in our system^21,22^.

### O_2_ microsensor measurements

Clark-type O_2_ microsensors (tip size =25 μm, response time < 0.2 s; OX-25 FAST, Unisense, Aarhus, Denmark) were used to characterize photosynthetic performance of the bionic corals. Net photosynthesis was measured via linear O_2_ profiles measured with O_2_ microsensors from the surface into the overlying diffusive boundary layer^2^. The sensors were operated via a motorized micromanipulator (Pyroscience, Germany). The diffusive O_2_ flux was calculated via Fick’s first law of diffusion for a water temperature = 25°C and salinity = 30 using a molecular diffusion coefficient for O_2_ = 2.255 × 10^-5^ cm^2^ s^-1(2)^. Gross photosynthesis was estimated via the light-dark shift method^41^. A flow chamber set-up provided slow laminar flow (flow rate = 0.5 cm s^-1^) and a fiber-optic halogen lamp (Schott KL2500, Schott, Germany) provided white light at defined levels of incident irradiance (400-700 nm) (0, 110, 220, and 1200 μmol photons m^-2^ s^-1^)^2^. Photosynthesis-irradiance curves were fitted to an exponential function^42^.

### Fiber-optic microsensors

The fluence rate (= scalar irradiance), E0, within the bionic coral was measured using fiber-optic scalar irradiance microsensors with a tip size of 60-80 μm and an isotropic angular response to incident light of ±5% (Zenzor, Denmark). Fluence rate measurements were performed through the tissue at a vertical step size of 100 μm using an automated microsensor profiler set-up as described previously^2^. Fluence rate was normalized to the incident downwelling irradiance, Ed, measured with the scalar irradiance sensor placed over a black light well at identical distance and placement in the light field as the surface of bioprinted constructs.

### Scanning electron microscopy (SEM)

SEM images were taken with a Zeiss Sigma 500 scanning electron microscope. Samples were prepared in two different ways. To image the bionic coral skeleton made of PEGDA, samples were dried at room temperature and sputter coated with iridium (Emitech K575X Sputter Coater). To image the bionic coral tissue made of GelMA, samples were snap frozen with liquid nitrogen, and were then lyophilized in a freeze dryer (Freezone, Labonco) for 3 days. The overall shape could not be maintained, but microscale structures (such as micropores of GelMA) were well preserved. The samples were sputter coated with iridium (Emitech K575X Sputter Coater) prior to imaging on the SEM.

### Confocal laser scanning microscopy (CLSM)

To characterize microalgal aggregate size and distribution in 3D, a confocal laser scan microscope was used (Nikon Eclipse TE-2000U). Bionic corals were placed on a cover glass and imaged from below with a 641 nm laser. Confocal stacks of chlorophyll *a* fluorescence were acquired using a pinhole size of 1.2 μm, a vertical step size for z-stacking = 1 μm, and a x,y resolution of 0.6 μm. Particle segmentation and visualization of the data was performed in ImageJ and the NIS confocal elements software (Nikon). Particle segmentation was performed via manual thresholding of 229-4095 gray scale values, with a cleaning factor of 6x (this eliminates smaller particles that are not aggregates), hole filling and a smoothing factor of 2x. The segmented particles were analysed for surface area, volume and particle density per volume.

---

**Extended data Figure 1.**
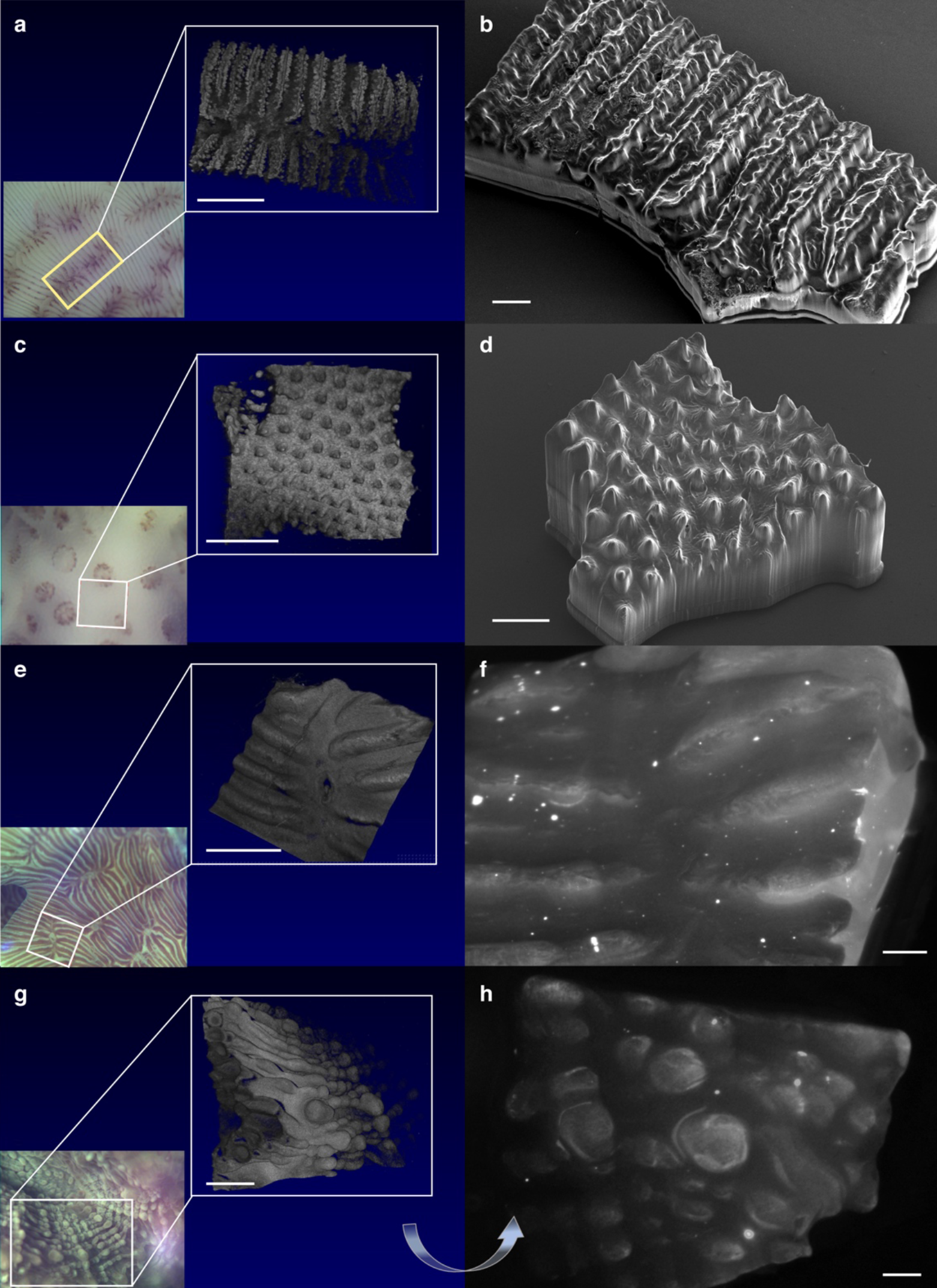
Microtopography of corals and 3D bioprinted bionic corals. Skeleton of *Pavona cactus* (**a, b**) and *Pocillopora damicornis* (**c, d**) as well as tissue surface of *Pavona cactus* (**e, f**) and *Favites flexuosa* (**g, h**). USB camera images and respective optical coherence tomography scans of natural corals (**a, c, e, g**) and 3D printed replica (**b, d, f, h**). Skeletal 3D printed constructs were imaged with an environmental SEM, while 3D printed tissue constructs were photographed with a microscope camera. Scale bar = 1mm (**a, b, d, e, g**) and 500 μm (**c, f, h**).

**Extended data Figure 2.**
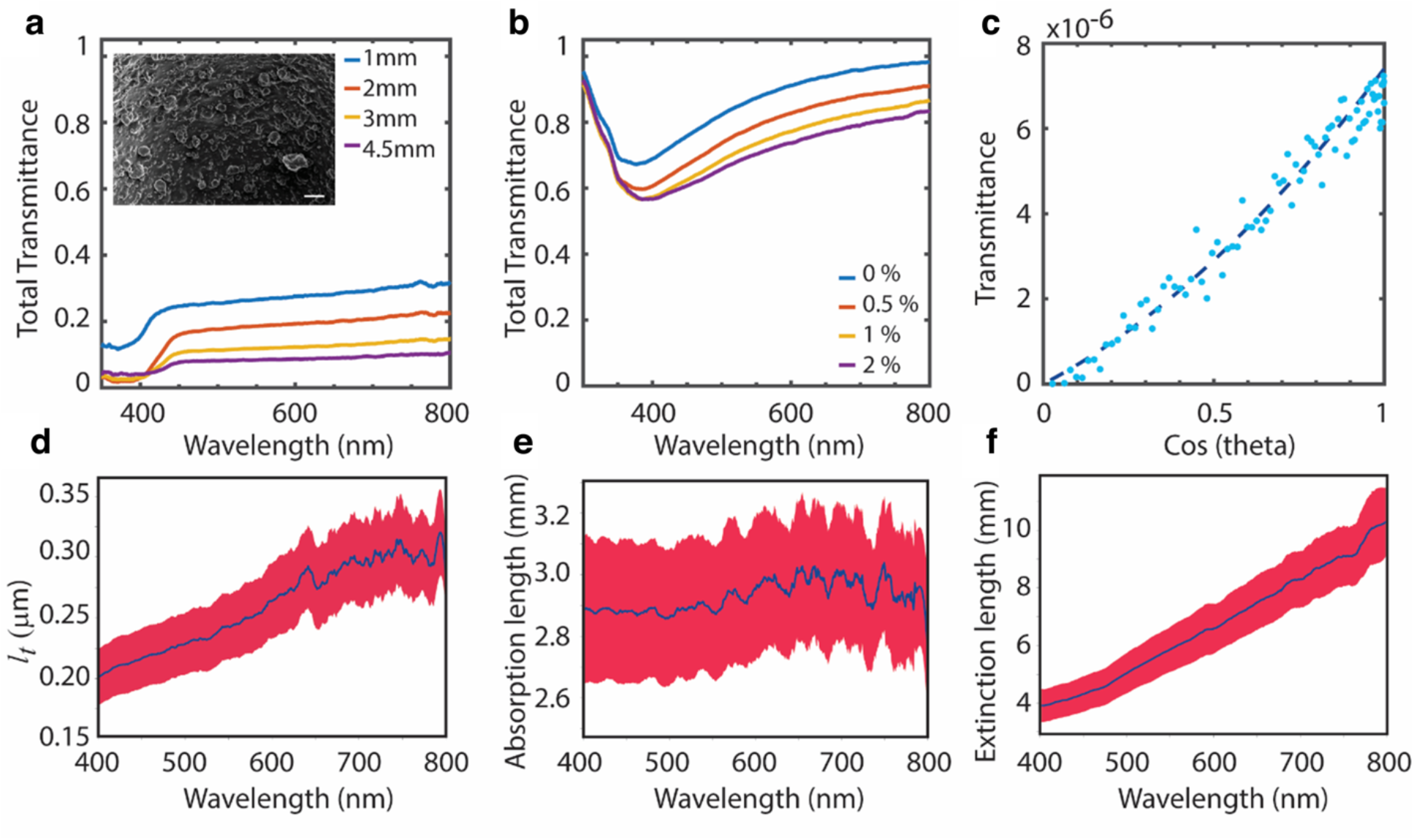
Optical characterization of 3D printed constructs. Total transmittance of bionic skeleton with 7% CNC concentration for different slab thicknesses (1-4.5mm) (**a**). The high CNC density yields a rough surface (see SEM image in inlet, scale bar = 40 μm). Total transmittance of bionic coral tissue doped with different concentrations of CNC (0-2%) (**b**). Fitting of extrapolation length (*z_e_*) for bionic skeleton according to Eq. 2 based on the angular distribution of transmitted light (**c**). Calculated transport mean free path (*l_t_*, μm) (**d**) and absorption length (*l_a_*, mm) for bionic skeleton (mean ± CI) (**e**). Extinction length for bionic tissue estimated using Beer-Lambert law (mean ± CI) **(f).**

**Extended data Figure 3.**
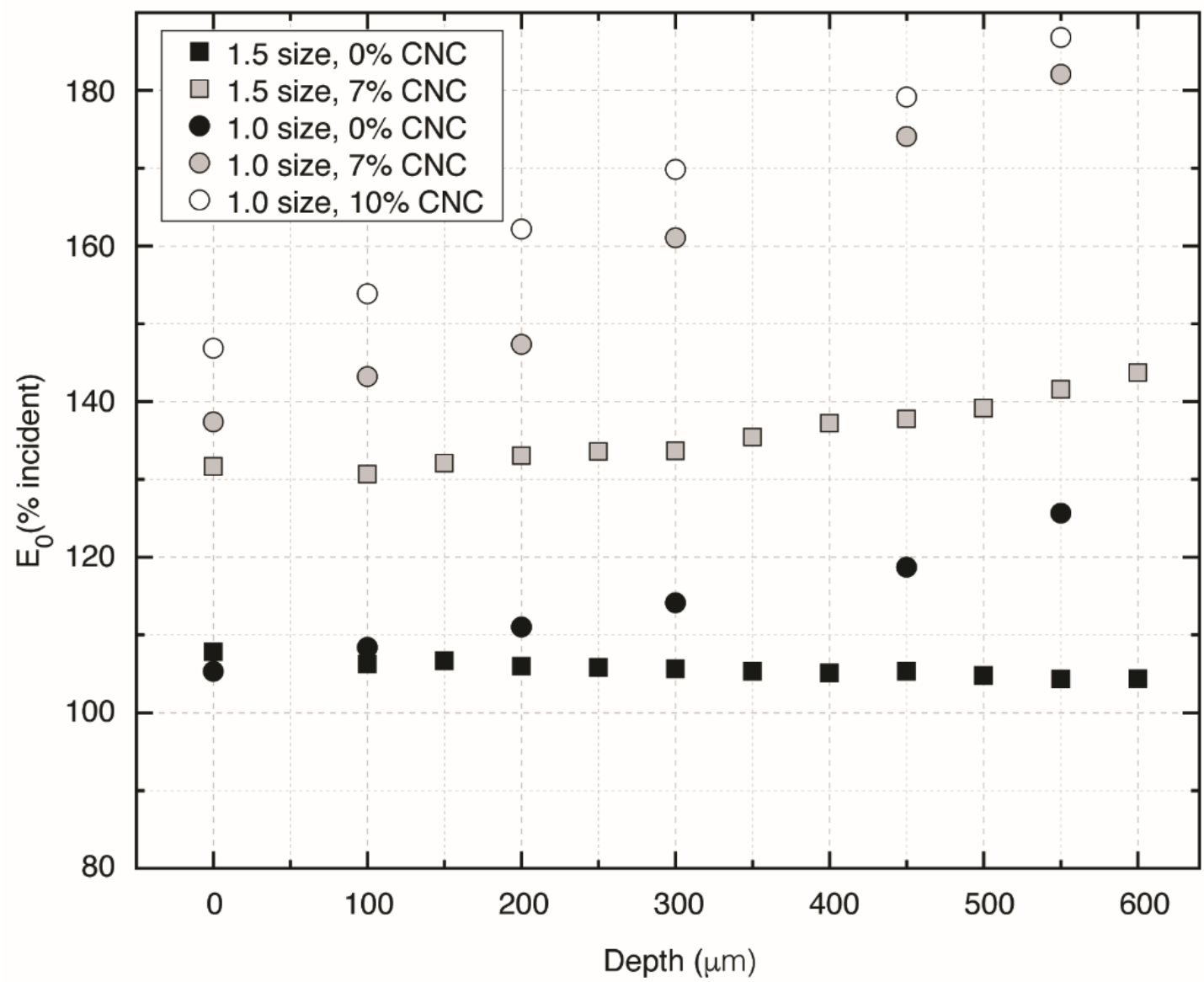
Effect of CNC doping and corallite cup size on fluence rate (E0) attenuation. Measurements were performed for different CNC concentrations (0-10%) using the original corallite cup size (maximal width = 1 mm) and a 1.5-fold enhanced size. E_0_ (fluence rate) was normalized to the vertically incident downwelling irradiance E_d_.

**Extended data Figure 4.**
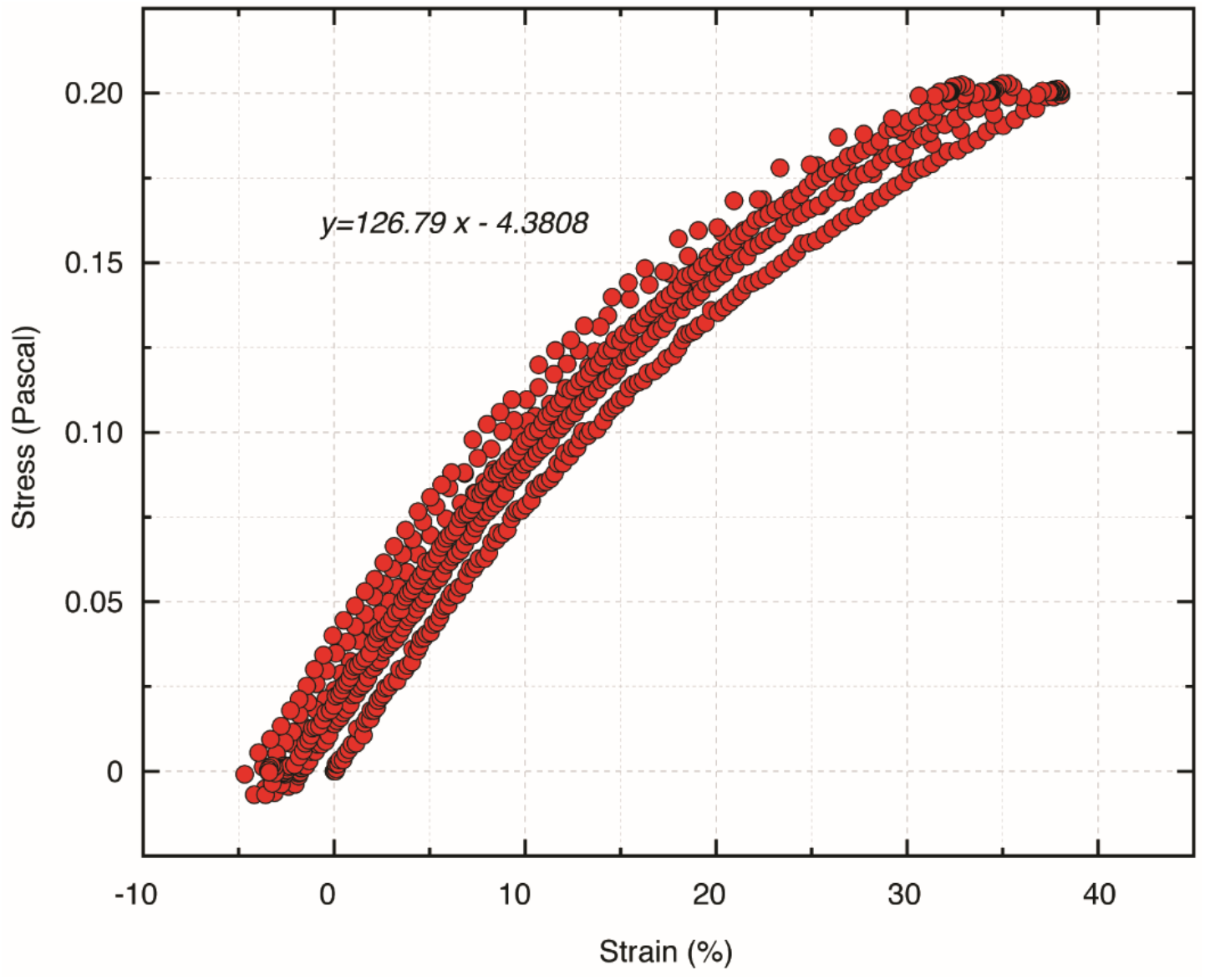
Stress-strain analysis of coral-inspired bionic tissue. Replicate measurements of 6 bionic tissues were performed. The average elastic modulus was *E* = 4.3 kPa.

## Acknowledgments

We thank Martin Tresguerres, Jennifer Smith, Qianqian Fang, Bryan Zhu, Debra Quick Jones, Haixu Shen, Jeffrey Alido, and Tressa Smalley for help with experimental and analytical work. Image credit in Fig. 1A is given to Gianfranco Rossi. Funding: This study was funded by the European Union’s Horizon 2020 research and innovation programme (702911-BioMIC-FUEL, DW), the European Research Council (ERC-2014-STG H2020 639088, SV), the David Phillips fellowship (SV), the National Institutes of Health (R21HD090662 and R01EB021857; SC), the National Science Foundation (1644967; SC), the Carlsberg Foundation (DW, MK), and the Villum Foundation (00023073; MK).

## Author contributions

conceptualized the study: DW, SV, DDD, MH, AS, MD; developed 3D printing approach: DW, SY, SC; developed optical model and characterized optical properties: DW, GI, SV; designed and performed cultivation experiments: DW, OG; performed imaging: DW, SY; provided materials: SC, FA, MK, SV, DDD; DW. All authors critically assessed the results and wrote the manuscript. Competing interests: The authors declare no competing interests. Data and materials availability: All data are available in the main text or the supplementary materials.

